# Revisiting the evolution of population stability due to selection for rapid development and early reproduction in *Drosophila*: the role of generation length

**DOI:** 10.64898/2026.09.04.749329

**Authors:** Medha Rao, Chinmay Temura, Amitabh Joshi

**Affiliations:** Evolutionary Biology Laboratory, Evolutionary and Organismal Biology Unit, Jawaharlal Nehru Centre for Advanced Scientific Research, Jakkur, Bengaluru 560 064, India

**Keywords:** density-dependent selection, population stability, generation length, population dynamics

## Abstract

The ubiquity of stable populations in nature generated considerable interest in how population stability might evolve, especially after the realization that higher per capita population growth rates typically yield unstable dynamics. Yet, most empirical and theoretical treatments implicitly assume that population dynamics and stability are invariant to generation length, an assumption that remains largely untested. Theory suggested that population stability could evolve as a correlated response to life-history evolution, which was first experimentally demonstrated using *D. melanogaster* populations selected for rapid development and early reproduction (FEJs). Constancy stability of FEJs evolved to be higher than their ancestral controls (JBs), likely due to correlated reductions in fecundity and pre-adult survivorship. However, that study assessed stability on a 21-day generation length (matching the JBs), resulting in a considerable mismatch with the generation length of FEJs (10 days). This raises a broader question of whether the observed differences in stability reflect evolved life-history changes or are an artifact of the generation length at which the populations were assayed.

To address this, we assessed the stability of FEJs, JBs, and relaxed-selection populations derived from the FEJs (CRFs and FRFs) across two generation lengths (12 and 18 days), tracking 320 small populations for 27 generations. Contrary to the earlier findings based on a 21-day generation length study, FEJs did not differ in constancy from JBs when assayed at a shorter generation length. Thus, even a modest difference in life-cycle length can significantly influence population stability, thereby underscoring the need to account for generation length when comparing stability across populations. Interestingly, constancy and persistence stability of selection regimes evolved in opposite directions, highlighting the need to assess stability along multiple axes. We discuss these results using an empirical framework, emphasizing its utility over simple population growth models for a richer understanding of population dynamics and stability.

## INTRODUCTION

Population stability is a central concern in population ecology and evolutionary biology (Dey & Joshi 2018). The stability of populations is closely tied to the factors that influence population growth, and traits that affect growth can, in turn, also shape stability (Mueller & Ayala 1981a). Population stability can be quantified in multiple ways (∼40 measures outlined in Grimm & Wissel 1997; Grimm *et al*. 1992), but the ecologically relevant metrics typically capture the ability of a population to remain unchanged, return to an ‘equilibrium state’ after a disturbance, or persist over time (Grimm & Wissel 1997). Among these, persistence is inversely related to a population’s extinction probability and therefore measures its ability to persist over time (Allen 1983). Constancy reflects the ability of a population to essentially remain unchanged and is typically assessed as the inverse of a measure of the degree of fluctuations around the average population size. Low constancy, characterized by large fluctuations around the mean population size, can also reduce the effective population size, potentially increasing the risk of genetic drift and the likelihood of extinction when population sizes are small.

Following the work of May (1974, 1976) and May and Oster (1976), a conundrum arose: simple population growth models demonstrated that an increase in the maximum per capita growth rate (*r)* beyond a threshold destabilizes the dynamics of the population. As *r* is a close correlate of fitness, all else being equal, natural selection should favor an increase in the intrinsic growth rate in populations and thus lead to more unstable populations. However, empirical studies showed that, contrary to expectations, natural and laboratory populations often exhibit relatively stable dynamics (Hassell *et al*. 1976; Thomas *et al*. 1980; Mueller & Ayala 1981b). Several hypotheses were proposed to explain this apparent paradox, including a set of ideas that consider the evolution of stability in populations. One class of theories posited that group-level selection could result in increased stability (Thomas *et al*. 1980; Berryman & Milstein 1989), because unstable local populations experience more frequent fluctuations in population size and often exhibit low densities, thereby increasing their susceptibility to extinction. Consequently, only stable populations would persist longer and hence, populations sampled in nature would tend to exhibit stable dynamics. However, the theory assumes that the differences in stability which lead to varying extinction rates among populations are heritable. If the environment were the primary factor driving variation in stability among populations, this proposed mechanism would not work (Mueller & Joshi, 2000).

Another class of theories proposed that selection on population-level demographic parameters, such as the growth rate (*r*) or the equilibrium population size (*K*), may enhance population stability (Hansen 1992; Ebenman *et al*. 1996). Mueller *et al*. (2000) tested this hypothesis using *Drosophila* populations maintained on food regimes that produced large, regular fluctuations in population size, but observed no evolutionary change in the constancy of the populations; persistence was not examined due to a lack of extinctions owing to very large population sizes. A third set of hypotheses suggested that stability could emerge as an indirect consequence of selection acting on individuals, thereby altering life-history traits that influence fitness. Therefore, stability could evolve as a by-product of life-history evolution, especially through trade-offs among life-history traits that impact population-level demographic parameters (Mueller & Joshi 2000). Various other explanations focused on ecological factors that could yield increased stability, and a lot of studies have examined both ecological and evolutionary explanations for population stability (reviewed in Mueller & Joshi 2000; Dey & Joshi 2018). These studies include both theoretical and experimental work, the latter largely done on laboratory populations of *Drosophila*, and have helped identify various factors that influence dynamics and stability, including larval rearing density, adult mortality, adult body size, female fecundity and its sensitivity to adult density, survivorship, sex-ratio, spatial variation in density, and constant number immigration (Mueller 1988; Mueller & Ayala 1981a; Sheeba & Joshi 1998; Mueller & Hyunh 1994; Sinha & Parthasarathy 1994; Jaggi & Joshi 2001; Tung *et al*. 2019). All these studies of dynamics in *Drosophila* populations typically treated generation length in discrete-generation experiments as a fixed aspect of the design, rather than as a variable that could influence estimates of population stability. Our present study examines, *inter alia*, the effect of generation time on population stability in the *Drosophila* system in which the evolution of population stability as a by-product of life-history evolution was first experimentally reported (Prasad *et al*. 2003; Dey *et al*. 2008).

Prasad *et al*. (2003) showed that *D*. *melanogaster* populations selected for rapid development and early reproduction (FEJs) also evolved higher constancy than their ancestral controls (JBs). The effect was primarily attributed to correlated reductions in fecundity and high larval mortality in the FEJs (Prasad 2003), both of which contributed to a lower amplitude of fluctuation in population size over time, relative to the JBs. A subsequent paper on a continuation of the study reported no significant difference in persistence between the FEJs and JBs (Dey *et al*. 2008), indicating that these two aspects of stability were not necessarily correlated.

Importantly, the studies of Prasad *et al*. (2003) and Dey *et al*. (2008) assessed population dynamics on a 21-day discrete generation length (the next generation was initiated every 21 days), corresponding to the generation length of the JBs during regular maintenance. FEJs, in contrast, were typically maintained on a 10-day generation cycle due to selection for rapid development and early reproduction. Thus, a 21-day generation length in the population dynamics experiment meant that the FEJ flies were well past the age at reproduction that they had been selected for, whereas the JBs were not. Over ∼125 generations of selection at the time, FEJs had evolved several correlated life-history changes, including reduced female fecundity, lower lipid reserves, smaller body size (Prasad 2003; Modak 2009), and shorter lifespan relative to the JBs (Modak 2009). Moreover, female fecundity in *Drosophila* decreases with age (Rose 1984). Given the shorter lifespan of FEJs, many individuals likely died before contributing to the next generation, at intervals of 21 days. Thus, age-linked effects on fecundity and survivorship may have shaped the observed dynamics in the FEJs, raising the possibility that their higher constancy stability was an artifact of the generation length used during the assay. To address this, we tested the effects of generation length on population dynamics across these selection regimes. We hypothesized that FEJs may not show greater stability relative to the JBs when assayed in a population dynamics experiment with a generation length closer to what they typically experienced during routine maintenance in their selection regime.

We also examined two sets of populations derived from the FEJs by relaxation of one or both selection pressures (FRFs and CRFs). The FRFs are relaxed only for rapid development, while the CRFs are relaxed for both rapid development and early reproduction. They are maintained at 12 day and 17 day generation lengths in the laboratory, respectively. These populations can aid in disentangling the contributions of rapid development and early reproduction, not only to life-history evolution, but also to population dynamics and stability. Our experiment tested the population stability of the four selection regimes (FEJ, JB, FRF and CRF) across two generation length treatments: 12 days and 18 days. The 12 day cycle approximates the native maintenance of the FEJs and FRFs, while the 18 day cycle is closer to the native maintenance of the JBs and CRFs.

Our results show that generation length plays a critical role in shaping population stability, and that observed patterns of relative stability among populations may not generalize across different generation lengths, underscoring the need to consider this often-overlooked aspect in investigations of population stability. Additionally, we show that constancy and persistence evolved in opposite directions across the various selection regimes, underscoring the need to assess population stability along multiple axes, as noted by Dey *et al*. (2008). The relaxed selection populations were found to have evolved various enhanced demographic attributes relative even to the JBs. We also used the realized population growth rate curves to explain patterns of population stability that conventional, simple population growth models would not capture (as in Pandey & Joshi 2026a,b).

## MATERIALS AND METHODS

### Experimental populations

We used sixteen large, outbred populations of *D*. *melanogaster*, all maintained at around 25°C under constant light and high humidity conditions. Four of these populations were selected for rapid development and early reproduction (Faster developing, Early reproducing JB-derived; FEJ_1-4_), while four served as the matched ancestral controls (Joshi Baseline; JB_1-4_). From the FEJs, two sets of relaxed selection populations were derived – one set of four populations was relaxed for the rapid development selection pressure alone (Faster development Relaxed FEJ-derived; FRF_1-4_), and the other set was relaxed for both rapid development and early reproduction (Completely Relaxed FEJ-derived; CRF_1-4_). The derivation and maintenance of these populations are described in detail elsewhere (For JBs: Sheeba *et al*. 1998; For FEJs: Prasad *et al*. 2000; For CRFs and FRFs: Mital 2019; Rao *et al*. 2025a). Replicate populations with the same numerical subscript share common ancestry (such as FEJ_1_, JB_1_, CRF_1_ and FRF_1_) and are, therefore, considered as randomized blocks in our statistical analyses. At the start of the dynamics assay, JBs had been maintained in the lab for 389 generations. FEJs had undergone 759 generations of forward selection. The FRFs and CRFs had undergone 199 and 135 generations of relaxed selection, respectively.

### Population dynamics experiment

We examined population dynamics under two discrete-generation lengths – 12 days and 18 days – using small, independent single-vial populations of *Drosophila*. Each population was initiated by introducing eight males and eight females into glass vials (2.2cm dia × 9.6 cm ht) containing ∼1.5 mL of banana-jaggery medium for 24 hours (denoted as Day 0). After 24 hours, the adults were discarded, and the eggs laid during this period formed the founder generation (i.e., Generation 1).

Emerging adults from each culture were collected daily and transferred to adult collection vials, which were maintained in parallel. These vials contained ∼4 mL of food medium, and fresh food was provided every other day. Flies were collected until day 10 and day 16 for the 12 and 18 day generation length treatments, respectively. Subsequently, adults were transferred to conditioning vials (see Food Regime section) for two days. On days 12 and 18 for the 12 and 18 day generation length treatments, respectively, surviving adults were transferred to fresh food vials containing ∼1.5 mL of food for a 24-hour oviposition period. Eggs laid in this window constituted the next generation (i.e., Generation 2). After the oviposition window, the adults were sexed, censused, and discarded. Thus, the small populations were maintained without any regulation on egg or adult densities for 27 generations (∼1.5 years), following which the experiment was terminated. The 12 day and 18 day generation cycles were run in parallel throughout the study.

For the 12 day generation cycle, flies eclosing beyond day 11 were excluded from the breeding pool, effectively truncating the eclosion distribution for some selection regimes (particularly the JBs). Flies in this treatment experienced the conditioning vials during days 10-12. In the 18 day discrete generation cycle, flies were collected from the culture vials till day 16. Any flies eclosing beyond this window were excluded from the breeding pool. Flies in this treatment experienced the conditioning vials during days 16-18.

### Food regime

Each small population was maintained under one of two food regimes: LL or LH. In both regimes, the larval food was limiting (exactly 1.5 mL per vial). In the LH regime, adults received ad libitum live yeast paste (smeared on the wall of the vial) during the two-day conditioning period, while adults in the LL regime did not. Both food regimes are known to destabilize and induce large fluctuations in population size over time, with the LH regime generating more substantial fluctuations (Dey 2012). Five replicate single-vial populations were established for each combination of selection regime, food regime, generation length, and block, resulting in a total of 320 small populations.

### Extinctions and restarting of small populations

An extinction event was recorded when no females were available to initiate the next generation. As the food regimes produced large fluctuations in population sizes, extinctions occurred frequently, and populations had to be restarted to continue the experiment. Extinct populations were reseeded with four females randomly sourced from the five replicate backup vials belonging to the same selection regime × generation length × block combination (run in parallel with the primary experimental vials). These females were provided 24 hours for oviposition, and the eggs laid initiated the next generation.

The backup vials were maintained at high larval food levels (exactly 6 mL), thereby reducing competition through high larval density during this stage. Adults were provided with ad libitum yeast during the conditioning period. Subsequent generations were initiated by randomly selecting only four females from the previous generation’s adult pool for the oviposition window, thereby imposing a density control even at the adult stage. Consequently, the backup vials comprised small, density-controlled populations, which were run in parallel with the assay vials. Five replicate backup vials were maintained for each combination of selection regime and block.

### Measurements of dynamics and stability

*Constancy* (Grimm & Wissel 1997) was quantified using the Fluctuation Index (FI; Dey & Joshi 2006), defined as:

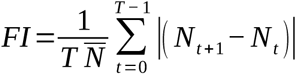

For each single vial population, *N_t_* is the population size in generation t, *N_t_*_+1_ is the population size in generation *t*+1, *T* is the total number of generations for which the assay was conducted and *N* is the average population size (across all generations). FI is inversely related to the constancy of the population.

*Persistence* (Grimm & Wissel 1997) was measured as the proportion of extinction events over the total number of generations for each population. Populations with a higher extinction probability were considered to have lower persistence stability. In earlier work, consecutive extinction events were treated as a single event to ensure the independence of each extinction, as females from other assay vials from whom eggs had already been harvested were used to reset any extinct vials (Dey *et al*. 2008). In our experimental design, as females from backup vials (run in parallel with the assay vials) were used to restart extinct populations, we considered all extinction events to be independent of one another. Nevertheless, we also looked at our data while treating consecutive extinctions as one extinction event (following Dey *et al*. (2008)), and found no qualitative differences in the results (Figure 2). Therefore, we primarily focus on the results from the analysis of total extinction probability. The extinction probability averaged over the five replicate populations for each combination of selection regime, food regime, generation length, and block was used as the unit of analysis.

### Measurements of demographic attributes

For each single vial population, three demographic attributes were estimated: the intrinsic per capita population growth rate (*r*), equilibrium population size (*K*), and the sensitivity of the realized population growth rate to density (*⍺*). The Ricker model (Ricker 1954) is known to effectively capture the dynamics of single-vial *Drosophila* populations (Sheeba & Joshi 1998), and is here parametrized as:

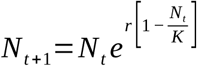

*N_t_* is the population size in generation *t*, *N_t_*_+1_ is the population size in the subsequent generation, *r* is the intrinsic growth rate, and *K* is the equilibrium population size.

The demographic estimates were obtained using the Levenberg-Marquardt method of nonlinear estimation in Statistica version 7 (Statsoft 1995). We also performed a semi-log transformation of the Ricker equation to get a linearized form. From this form of the equation, the demographic attributes were obtained by regressing Ln (*N_t+1_/N_t_*) on *N_t_*. In this case, the Y-intercept, X-intercept, and the slope were the estimates of *r*, *K*, and *⍺*, respectively. As both methods yielded qualitatively similar results, as has also been seen in earlier studies (e.g., Pandey & Joshi 2026a,b), we present the linear regression estimates in detail.

### Statistical analyses

Linear regressions were performed using the *lm* function in R (R Core Team 2025) to estimate *r*, *K*, and *⍺*. All graphs were plotted using R packages *tidyverse* and *ggplot2* (Wickham 2016; Wickham *et al*. 2019).

Estimates of the demographic attributes (*r*, *K,* and *⍺*), and population stability (constancy and persistence) were analyzed using mixed-model ANOVAs, with selection regime, generation length, and food regime as fixed factors and replicate blocks as a random factor in each analysis. Realized growth rates were analyzed using the aforementioned method, with population-size bins as an additional fixed factor. Post-hoc comparisons were performed using Tukey’s HSD. ANOVAs were performed in R release 4.5.1 (R Core Team 2025) using the *aov* function in the *stats* package (R Core Team 2025).

## RESULTS

### Effect of generation length on constancy

As expected from earlier studies (Dey 2007; Tung *et al*. 2019), the LH food regime induced significantly higher fluctuations in the populations, yielding FI values that were 39.4% higher than the LL regime (Figure 1; Table S1, main effect of food regime, *F*_2,3_ = 603.7, *P* < 0.001). This pattern held across both generation length treatments, but the difference was more pronounced in the 18 day cycle, with 87.3% higher FI in LH than in the LL regime. In the 12 day cycle, the LH regime had 46.9% higher FI than the LL regime (Figure 1; Table S1, interaction effect between food regime and cycle length, *F*_4,3_ = 19.42, *P* = 0.02). The FI values observed in the LH regime were consistent with those seen earlier (Dey 2007; Dey *et al*. 2008, 2012; Dey 2012; Tung *et al*. 2019; Pandey and Joshi 2026a,b), as were those in the LL regime (Dey 2007; Dey 2012; Tung *et al*. 2019).

**Figure 1.**
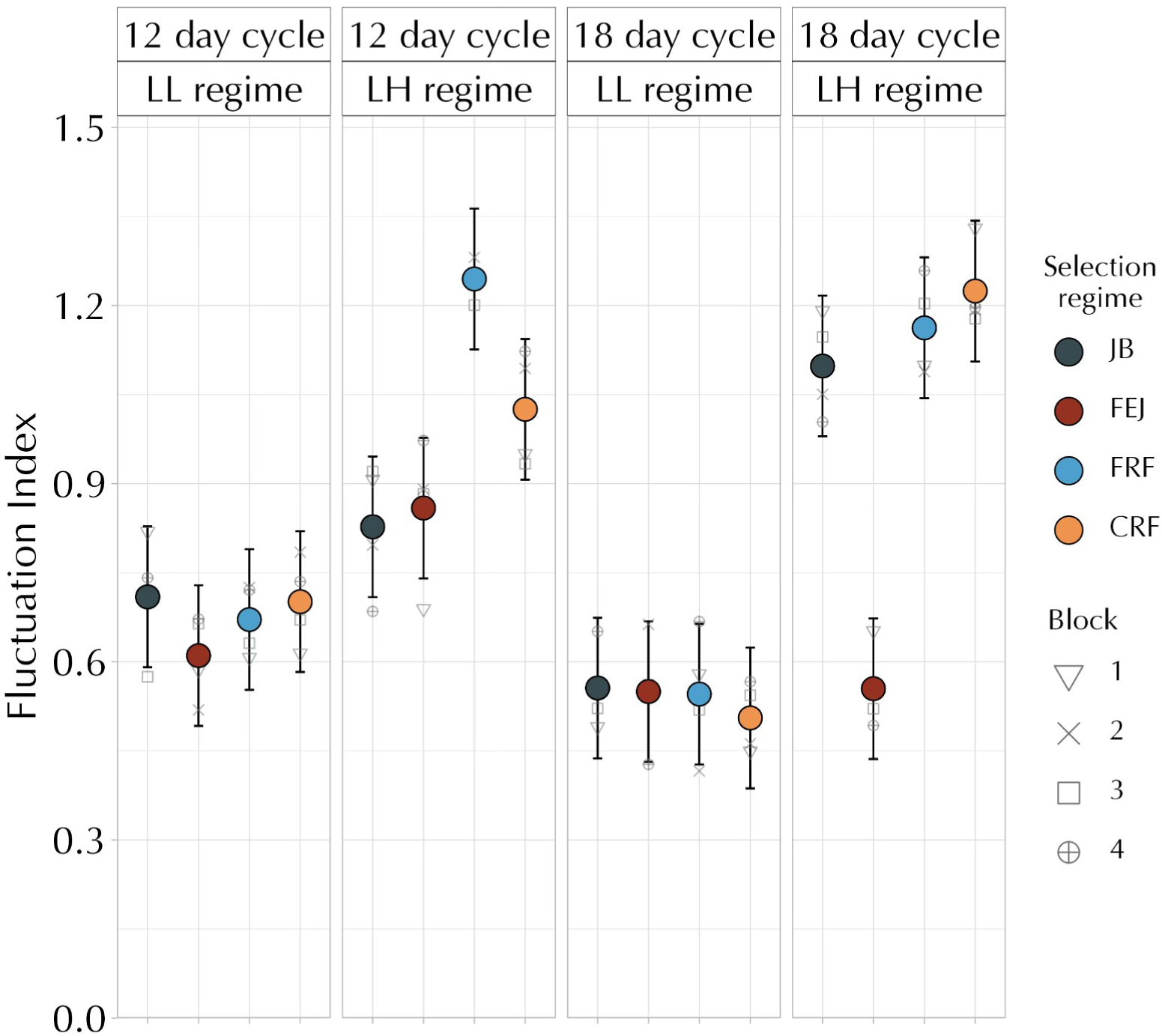
Mean fluctuation index (FI) of the selection regimes across all combinations of generation length and food regimes, when averaged over replicate blocks. Constancy is inversely related to the FI value. The error bars represent 95% confidence intervals and can be used for visual hypothesis testing. The lack of an overlap between the error bars of any two groups signifies a statistically significant difference between their respective means.

On average, FRFs and CRFs had the highest FI, followed by JBs and FEJs (Figure 1; Table S1, main effect of selection, *F*_3,9_ = 36.51, *P* < 0.001), albeit the magnitude of these differences was dependent on the exact combination of food regime and cycle length (Figure 1, Table S1, interaction effect between cycle length, food regime, and selection regime, *F*_3,9_ = 18.10, *P* < 0.001). Notably, the differences among selection regimes were limited to the LH food regime (Table S1, Interaction effect between selection regime and food regime, *F*_3,9_ = 22.14, *P* < 0.001). In the LL regime, on both 12 and 18 day cycles, FI values of all four selection regimes were very similar (Figure 1). In the LH regime on a 12 day cycle, FI values of CRFs and FRFs were substantially higher than those of the FEJs and JBs, which did not differ among themselves. However, in LH on an 18 day cycle, CRFs, FRFs and JBs showed similar values of FI, with all three being significantly higher than the FEJs (Figure 1, Table S1, interaction effect between cycle length, food regime, and selection regime, *F*_3,9_ = 18.10, *P* < 0.001). Incidentally, except for FEJs on an 18 day cycle, all FI values in the LH food regime are consistent with typical LH FI values shown by a variety of populations (Dey *et al*. 2008, 2012; Dey 2007, 2012; Tung *et al*. 2019; Pandey and Joshi 2026a,b).

The main takeaway from this set of results is that (a) reducing the generation length in the population dynamics assay to one more commensurate with the normal FEJ maintenance schedule can render the constancy of FEJs and JBs similar, unlike what was seen earlier with a JB-like generation length (Prasad *et al*. 2003; Dey *et al*. 2008), and (b) the CRFs and FRFS, by and large, have reduced constancy compared to the JBs.

### Effect of generation length on persistence

As expected from earlier studies (Dey 2007; Dey & Joshi 2013), the LH food regime, on average, resulted in a higher number of extinctions than the LL regime (Figure 2a, Table S2, main effect of food regime, *F*_1,3_ = 33.36, *P* = 0.010). This was true for FRFs, CRFs, and JBs (Figures 2a & S1, Table S2, interaction between selection regime and food regime, *F*_3,9_ = 9.020, *P* < 0.01). Interestingly, FEJs underwent fewer extinctions in the LH treatments than the LL treatment, even though the LH food regime is known to reduce persistence (Dey 2007, 2012; Tung *et al*. 2019; Figures 2a & S2a). However, this was entirely due to the 18 day cycle results: FEJs underwent more extinctions in LL than LH only on an 18 day cycle, not on the 12 day cycle (Figure S1). On average, the LH food regime caused significantly higher levels of extinction in the 12 day cycle length treatments relative to the 18 day treatments. In comparison, the LL food regime, on average, resulted in very similar levels of extinction across both cycle length treatments (Figure 2, Table S2, interaction between food regime and cycle length, *F*_3,3_ = 21.635, *P* = 0.018). The lack of a significant difference in this case could be due to the averaging out of effects in these treatment combinations caused by the contrasting behavior shown by FEJs and the rest of the selection regimes. FEJs had a significantly higher number of extinctions in 18 day cycle treatments compared to other selection regimes, while the other selection regimes showed somewhat higher extinction probability in the 12 day cycle length treatment (Figures 2, Figure S2b), a pattern perhaps due to their early life fecundity being substantially higher than that of the FEJs.

**Figure 2.**
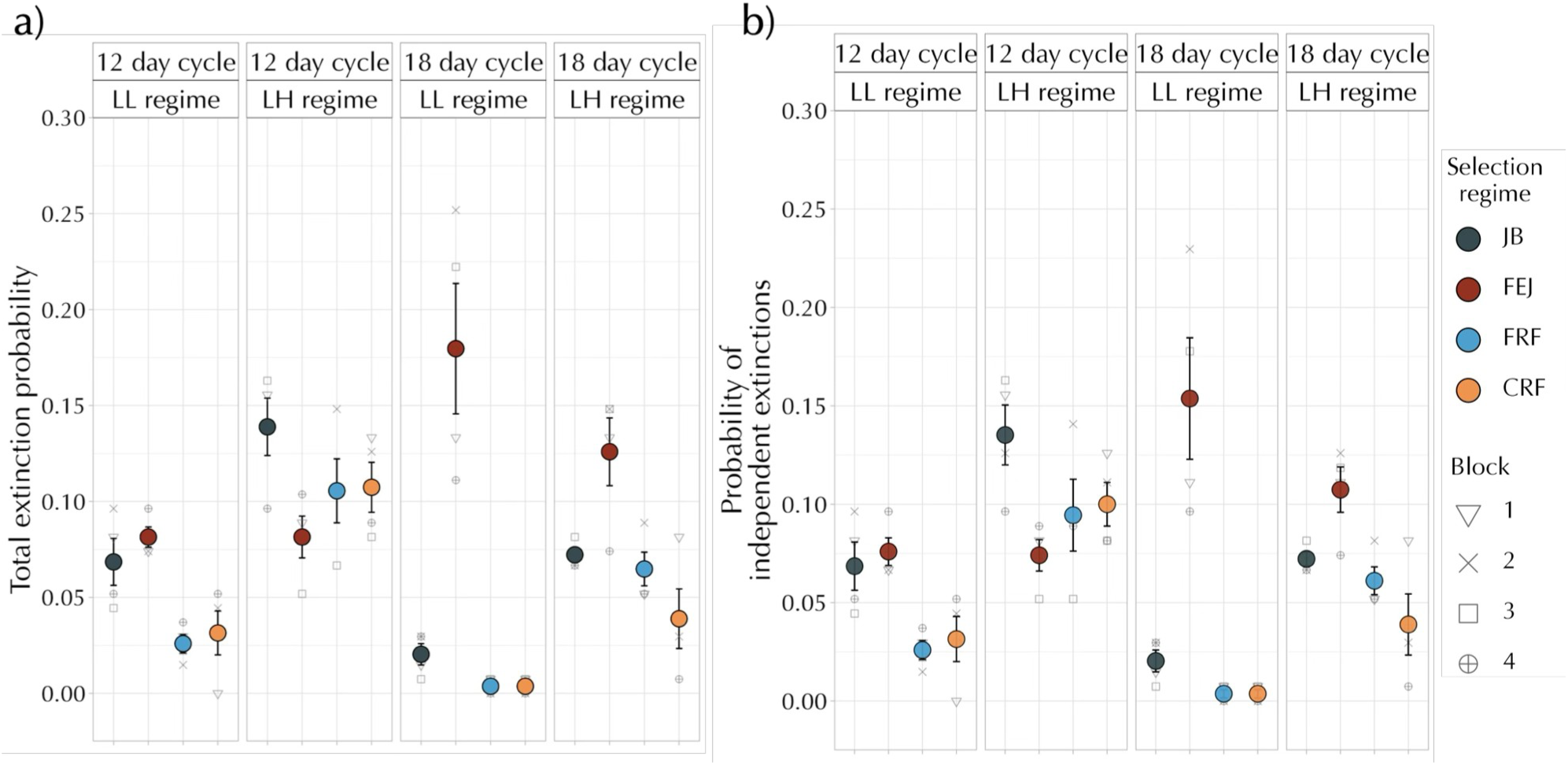
**a)** The average probability of extinction (per generation), and **b)** the average probability of independent extinctions (per generation), considering consecutive extinctions as one extinction event, of the selection regimes across two generation lengths and food regimes. Persistence stability is inversely related to extinction probability. The error bars represent the standard error around the mean.

On average, among selection regimes, FEJs showed the highest probability of extinction, followed by the JBs, whereas the CRFs and FRFs exhibited markedly lower extinction rates (Table S2, main effect of selection regime, *F*_3,9_ = 49.36, *P* < 0.001). Broadly, these qualitative patterns were consistent across treatments (Figure 2). The only exception to this is in the case of a 12 day cycle length and LH treatment combination, where FEJs showed relatively the lowest extinction probability (Figure 2). Importantly, these results demonstrate two aspects regarding the evolution of persistence. First, that FEJs had higher persistence than JBs in the 12 day, LH combination (∼5.75% higher), but a lower persistence than the JBs in the 18 day, LH treatment (∼5.38% lower) (this treatment was closest to the previous study), suggesting a significant role of generation length in shaping persistence. Second, CRFs and FRFs consistently showed lower extinction probabilities than JBs, particularly in the 12 day treatments, indicating that the relaxed selection populations had evolved higher persistence than even their ancestral populations before the forward selection (Figures 2 & S2b).

No qualitative differences were observed between the two methods of estimating extinction probability – either by considering all extinction events to be independent or by considering consecutive extinctions as a single extinction (compare Figure 2a and 2b). The only considerable difference between the two methods of estimation is the total number of extinctions experienced by the FEJs, particularly in the 18 day treatment (Figure 2), indicating that consecutive extinctions were primarily experienced by the FEJs, and that too in this treatment condition.

### Relaxed-selection populations maintained higher equilibrium population sizes (K) than the ancestral and forward-selected populations

The CRFs and FRFs maintained significantly higher equilibrium population sizes (*K*) than both the JBs and FEJs (Table S3, main effect of selection, *F*_3,9_ = 59.4, *P* < 0.001). This pattern remained consistent across the cycle length treatments, albeit to varying degrees (Table S3, interaction effect between selection and cycle length, *F*_3,9_ = 12.6, *P* < 0.01). The extent of divergence between the relaxed-selection populations from both the forward-selected and ancestral populations was the highest in the LL food regimes, across both types of generation lengths (Figure 3a, Table S3, interaction effect between selection, cycle length, and food regime, *F*_3,9_ = 11.91, *P* < 0.01).

**Figure 3.**
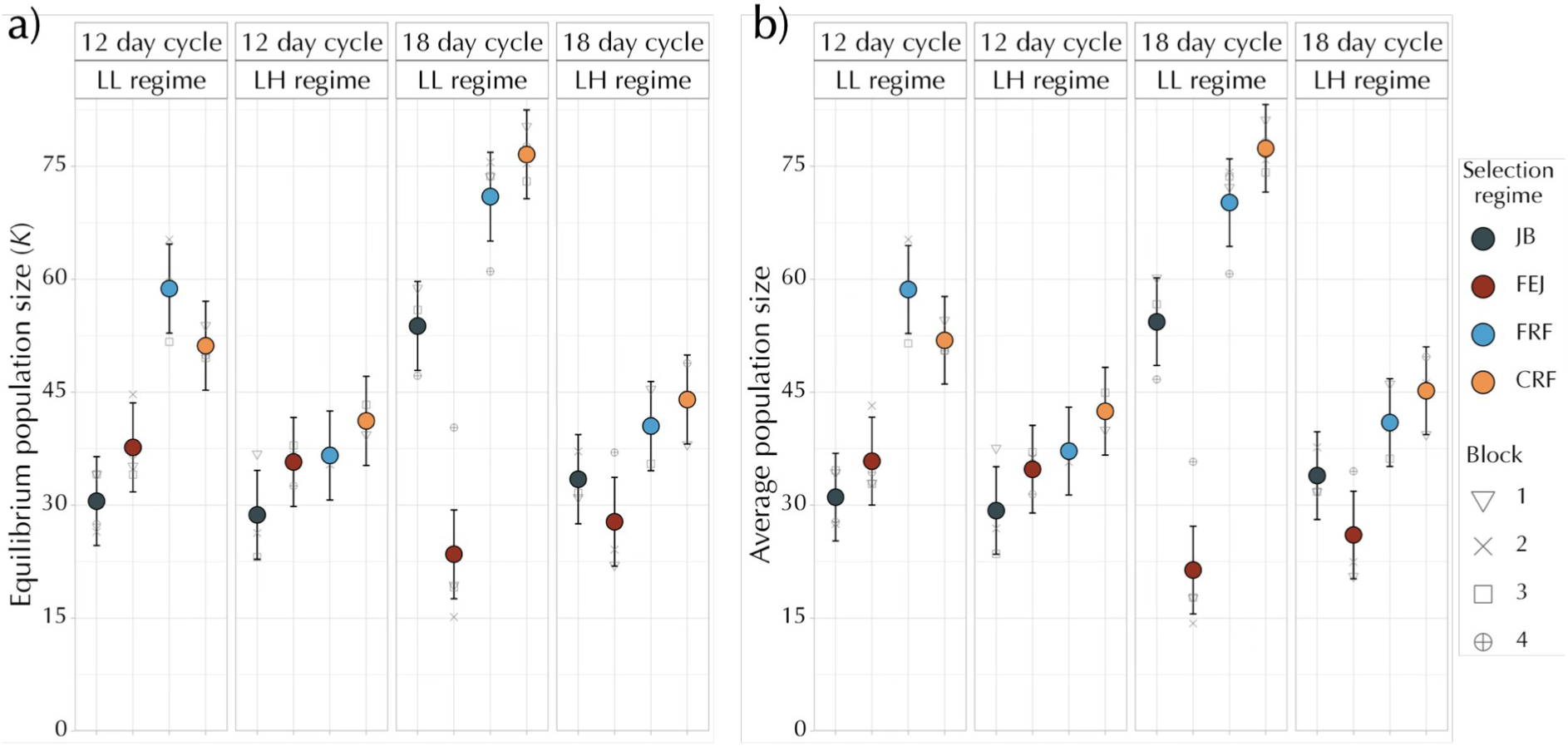
**a)** Estimates of the mean equilibrium population size (*K*), and **b)** the average population size attained by the different selection regimes over the course of the dynamics assay, across all treatment combinations when averaged over replicate blocks. The error bars represent 95% confidence intervals and can be used for visual hypothesis testing. The lack of an overlap between the error bars of any two groups signifies a statistically significant difference between their respective means.

In the 18 day cycle treatment, under the LL food regime, FEJs attained significantly lower *K* than all other selection regimes (Figure 3a). In all other treatment combinations, FEJs and JBs did not differ significantly from each other. However, it should be noted that in terms of the pattern, FEJs attained lower *K* than JBs in the 18 day treatments (56.3% lower in LL and 17.01% lower in LH, respectively), while the reverse occurred in the 12 day treatments (23.6% higher in LL and 24.3% higher in LH, respectively), indicating a possible impact of generation length yet again (Figure 3a). The patterns in the estimates of *K* closely matched those in the average population size obtained from the time-series data, as shown in Figure 3b; this indicates that simple models of population growth are likely to capture the dynamics reasonably well.

### Relaxed-selection populations exhibited higher intrinsic growth rates (r) than the ancestral and forward-selected populations

Overall, and consistent with previous studies (Dey 2007; Dey 2012), populations subjected to the LL food regime showed significantly lower intrinsic growth rate (*r*) estimates than those kept in the LH regime, primarily in the 18 day cycle length (Figure 4, Table S4, main effect of food regime, *F*_1,3_ = 30.8, *P* = 0.011). Populations also had, on average, lower estimates of *r* in the 18 day cycle treatment than in the 12 day treatment (Table S4, main effect of cycle length, *F*_1,3_ = 24.72, *P* = 0.015). However, this result seems driven by the difference between mean estimated *r* between LL 12 day and LH 18 day (Figure 4), and might not have great biological relevance, especially given the very similar realized population growth rates across densities exhibited by the CRFs, FRFs and JBs across the 18 day and 12 day treatments (Figure 5).

**Figure 4.**
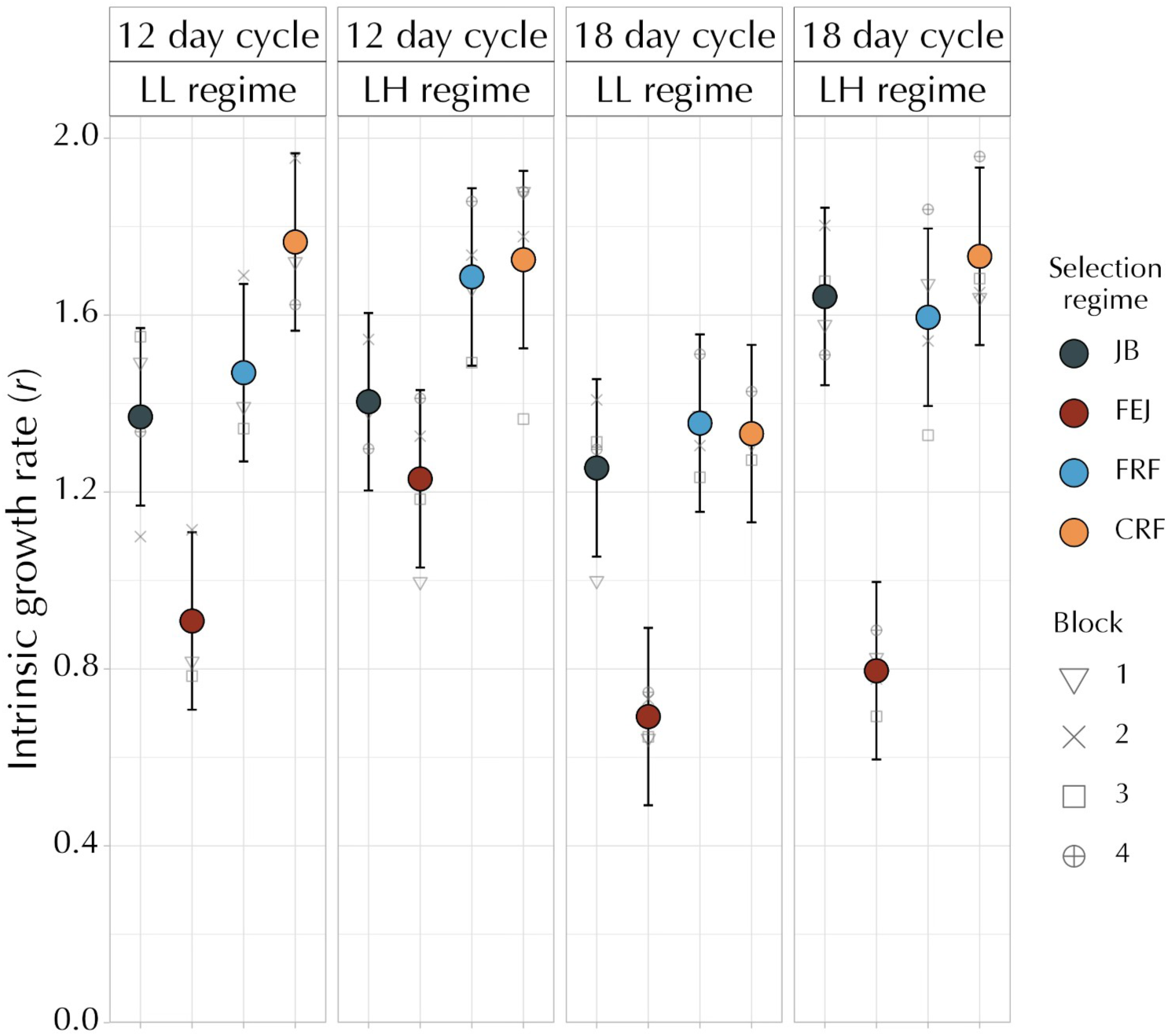
Estimates of the mean intrinsic growth rate (*r*) of selection regimes across all treatment combinations over the course of the dynamics assay, averaged over replicate blocks. The error bars represent 95% confidence intervals and can be used for visual hypothesis testing. The lack of an overlap between the error bars of any two groups signifies a statistically significant difference between their respective means.

**Figure 5.**
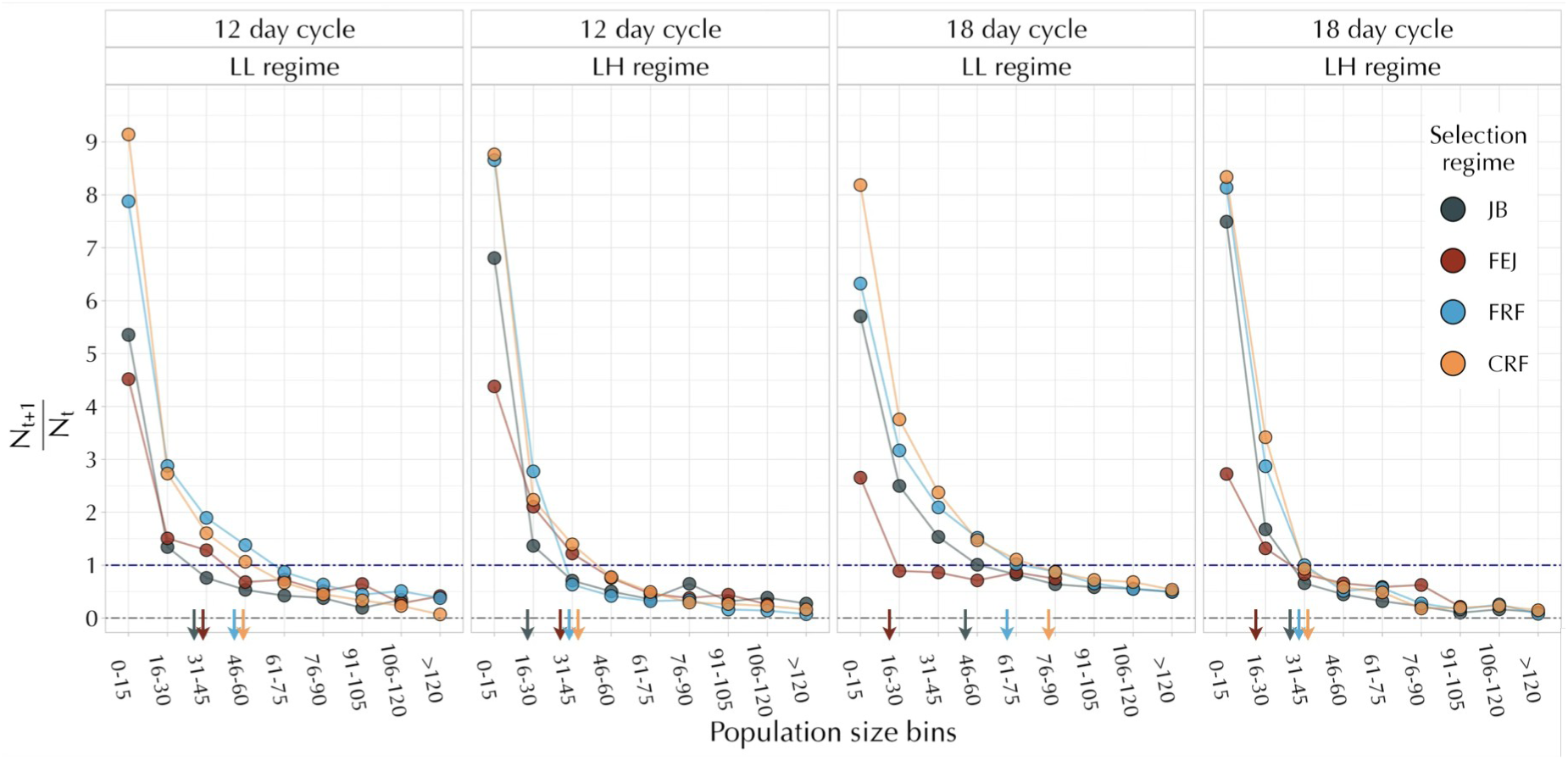
Mean realized growth rates 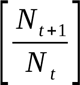 of the selection regimes across all treatment combinations and population size bins, when averaged over all occurrences belonging to a particular bin for all replicate populations. Population sizes from 0 to 120 were categorized across bins of 15 individuals. Population sizes beyond 120 individuals were pooled into one bin, labeled as > 120. The arrows indicate the population size bin where the equilibrium population size (*K*) for each treatment combination lies. The arrows are color-coded to match the selection regime they represent.

FRFs and CRFs maintained higher values of *r* than the JBs and FEJs on average, but the pattern was restricted to the 12 day cycle length (Figure 4, Table S4, main effect of selection, *F*_3,9_ = 63.2, *P* < 0.001). FEJs consistently had the lowest *r* estimates among all selection regimes across all the treatments. Only in the 12 day LH treatment combination, was the *r* estimate of FEJs not significantly lower than the JBs (Figure 4, Table S4, interaction between selection, cycle length, and food regime *F*_3,9_ = 5.809, *P* = 0.017). On average, the FEJ-JB differences were more prominent in the 18 day cycle treatments (Figure 4, Table S4, interaction between selection and cycle length, *F*_3,9_ = 4.06, *P* = 0.04).

For the JBs, CRFs and FRFs, differences in *r* between the LL-LH food regimes are, on average, greater for the 18 day treatment (∼30%, ∼30.1% and ∼17.6% increase in *r* in the LH treatment, respectively) compared to the 12 day treatment (∼2.5%, ∼2.2% and ∼14% increase in *r* in the LH treatment; Figure 4). This pattern likely reflects an interaction between adult yeasting, age-dependent decline in fecundity, along with differences in the basal level of fecundity among the selection regimes (*i.e.*, JB > CRF > FRF > FEJ; see Rao *et al*. 2025a).

JBs and CRFs are known to show high levels of basal fecundity in the 12 day treatment (Rao *et al*. 2025a), which possibly does not increase beyond a point, even with the presence of yeast. Consequently, even under the LL regime, they maintain high growth rates, leading to a lowered difference in *r* between the food regimes in the 12-day treatment. In contrast, in the 18 day treatment, as flies are substantially older, fecundity declines, resulting in lower *r* values for the LL regime. In this scenario, the provision of yeast in the LH regime can result in much higher fecundity for these populations, resulting in a greater difference between the LL-LH regimes. FRFs show a comparatively smaller difference between the LL-LH regimes across both cycle length treatments. This is likely due to their lower basal fecundity relative to the JBs and CRFs. The proportional increase in fecundity after yeasting may be more moderate relative to the JBs and CRFs, leading to the observed patterns.

These observed patterns are also consistent with the FEJs showing a large difference in growth rates between the LL-LH regimes, even in the 12 day cycle (∼35.3% increase in *r* in the LH treatment), as FEJ fecundity is very low even early in life. FEJs experience a pronounced age-related drop in fecundity by the relevant window for the 18 day treatment (see Rao *et al*. 2025a), which potentially cannot be rescued by the provision of yeast, leading to a relatively lower difference in *r* between the LL and LH regimes (∼14.9% increase in *r* in the LH treatment; Figure 4).

### Relaxed-selection populations showed a lower sensitivity to increasing density (⍺) than ancestral populations

Overall, the estimates of *⍺* (Figure S3) were similar, though slightly higher, to those seen in another study of this kind (Pandey and Joshi 2026a,b). Overall, the JBs showed the greatest sensitivity of realized population growth rate to density (most negative *⍺*) in 3 out of 4 combinations of cycle length and food regime, whereas *⍺* values of FEJs were similar to the CRFs and FRFs in 3 of the four combinations (Table S5, Figure S3). Another general pattern was that the relaxed-selection populations generally showed a less negative value of *⍺* than JBs, suggesting they were less sensitive to density increases (Figure S3). Moreover, the absolute differences in *⍺* between the CRFs or FRFs and JBs were almost twice as large compared to those previously seen between populations adapted to larval crowding (with reduced sensitivity to density) and their ancestral controls (Pandey & Joshi 2026a,b).

The picture becomes clearer on examination of the realized population growth rate versus density curves across all treatment combinations. These curves provide insight into how each selection regime responded to changes in adult density (Figure 5). The y-axis 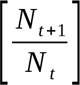 represents the average realized growth rate of replicate populations within the selection regime in the four combinations of cycle length and food regime. Values >1 imply increased population size in the next generation, values < 1 imply a decline in the population size, and a value of 1 indicates that the population exactly replaced itself in the next generation.

Across all treatment combinations, FEJs consistently had the lowest realized population growth rate at low adult density (i.e., in the bin of 0-15 individuals) compared to the other selection regimes (Figure 5). The relaxed selection populations (FRFs and CRFs) showed higher growth rates in low-density bins than FEJs and JBs. While all selection regimes exhibited a steep decline in growth rate when the density increased to 16-30 individuals, the relaxed populations still showed a higher growth rate compared to JBs and FEJs (Figure 5).

Notably, their growth curves remained above 1 for a broader range of population size bins (relative to the JBs or the FEJs), indicating their ability to maintain (or increase) their population sizes over this range. In the LL regime in particular, the realized growth rate of FRFs and CRFs remained ≥ 1 until the population size crossed 60 individuals (Figure 5). In contrast, while the JBs displayed a growth rate higher than the FEJs at low density, it remained consistently lower compared to the FRFs and CRFs. Additionally, JBs fell below the growth rate of 1 much sooner compared to FRFs and CRFs, particularly in the LL regime, indicating a more negative effect of increasing density on the growth rate of JBs (Figures 5 & S3). These patterns would, in part, likely be due to the generally higher *K* values in the CRFs and FRFs across most regimes (Figure 3a).

We performed an ANOVA for the realized growth rate with population size bins as an additional fixed factor for all selection regimes. For this analysis, we only considered bins that had representation across all four replicate blocks (i.e., 0-15, 16-30, 31-45, and 46-60). On average, FEJs had a significantly lower realized growth rate than the relaxed populations, while JBs had intermediate values (Figure 6b, Table S6, main effect of selection regime, *F*_3,9_ = 26.96, *P* < 0.001). The lowest bin, corresponding to 0-15 individuals, had the highest realized growth rate (Table S6, main effect of population size bins, *F*_3,9_ = 210.08, *P* < 0.001). Both FEJs and JBs did not differ significantly from a realized growth rate of 1 in the second bin (of 16-30 individuals). For the same bin, CRFs and FRFs exhibited a growth rate significantly higher than 1 (Figure 6a, Table S6, interaction between selection regime and population size bins). Additionally, at the second-highest bin of 31 to 45 individuals, the relaxed selection populations had a growth rate significantly higher than zero. In contrast, growth rates of JBs and FEJs did not statistically differ from zero (Figure 6a).

**Figure 6.**
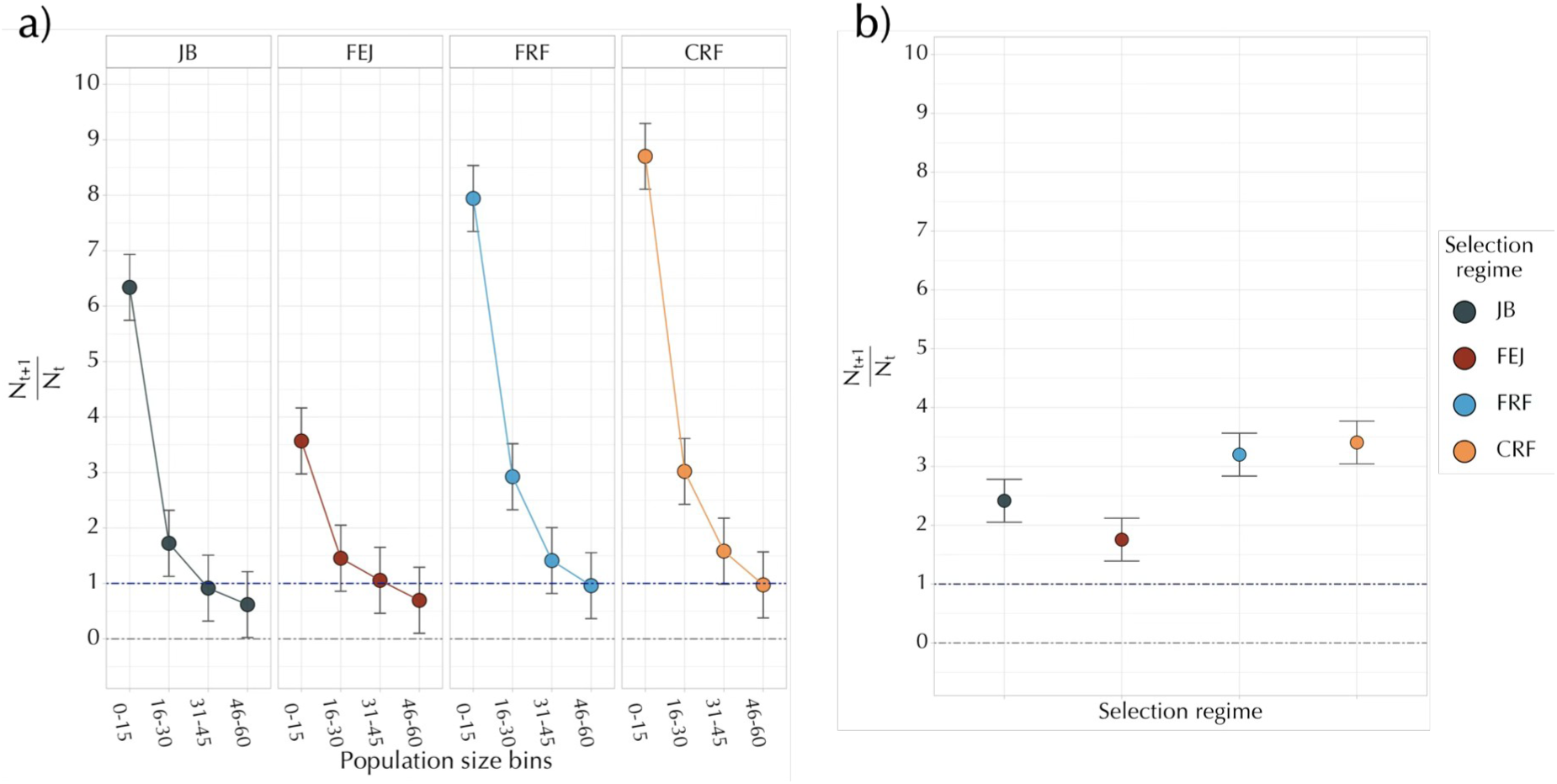
Mean realized growth rate 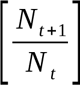 across all treatment combinations of a) selection regimes in different population size bins and b) selection regimes, when averaged over other factors. In this analysis, only bins that had a representation from each replicate block were considered (0-15, 16-30, 31-45, 46-60). The error bars represent 95% confidence intervals and can be used for visual hypothesis testing. The lack of an overlap between the error bars of any two groups signifies a statistically significant difference between their respective means.

## DISCUSSION

Earlier studies on the FEJs and JBs found that selection for rapid development and early reproduction led to the correlated evolution of increased constancy, but not persistence (Prasad *et al*. 2003; Dey *et al*. 2008). However, those studies were conducted on a 21-day generation cycle, which corresponded to the routine maintenance of the JBs, but resulted in a substantial mismatch for the FEJs, which were routinely maintained on a 10-day generation cycle as part of their selection regime. This mismatch raised the possibility that the observed stability differences between the JBs and FEJs might be a consequence of assay conditions and thus represent a strong genotype × environment interaction rather than a straightforward correlated evolution of greater constancy in the FEJS, as it was interpreted at that time. Our results clearly show that generation length in the actual population dynamics assay does have a significant impact on constancy and persistence in a selection regime specific manner, and conclusions drawn from mismatched assay conditions may not hold when populations are assayed closer to their native maintenance conditions. This has important implications for the design of population dynamics experiments in which populations differ in their ‘native’ generation length.

Another aim of the present study was to examine the population dynamics and stability characteristics of the relaxed selection FRF and CRF populations in order to ascertain the relative contribution of the forward selection pressures on development time versus timing of reproduction to the earlier observed differences (Prasad *et al*. 2003; Dey *et al*. 2008) in stability between the FEJs and JBs., and to assess how similar to the JBs the relaxed selection populations had become with regard to dynamics and stability characteristics. Overall, there were no consistent differences between CRFs and FRFs for any of the attributes examined: constancy (Figure 1), persistence (Figure 2), equilibrium population size (Figure 3), intrinsic population growth rate (Figure 4), sensitivity of realized growth rate to density (Figures 5, S3), or the degree to which LL and LH regimes induced sharp two-point cycles (Figure S4). This could be taken as an indication that perhaps the selection for rapid development played a much bigger role than selection for early reproduction in mediating the evolution of differences observed by Prasad *et al*. (2003) and Dey *et al*. (2008) between the FEJs and JBs in terms of population dynamics and stability attributes, since both the FRFs and CRFs were relaxed for selection on development time, but only the CRFs were also relaxed for selection on age at effective reproduction. However, we must also consider the fact that the FRFs had undergone 47% more generations of relaxed selection compared to the CRFs (199 versus 135 generations) at the time of the population dynamics experiment reported here. Thus, it is possible that CRFs, over an equal number of generations of relaxed selection as the FRFs, might have diverged from the latter, thus reflecting a substantial role for selection on early reproduction in explaining the earlier observed differences in stability between the FEJs and JBs. Surprisingly, in comparison to the JBs, both sets of relaxed selection populations seemed to have become slightly less stable – especially in constancy (Figure 1), equilibrium population size (Figure 3), and the propensity to exhibit sharp two-point cycles (Figure S4) – indicating that relaxed selection had resulted in these populations exceeding trait values of the ancestors of the original forward selected populations. While this can happen in relaxed selection scenarios (Teotónio & Rose 2001; Teotónio *et al*. 2002), we cannot rule out the possibility that the trait values of the JBs (and of the FEJs) have changed in the few hundred generations between the present study and the earlier studies of Prasad *et al*. (2003) and Dey *et al*. (2008). We next discuss some aspects of our results in greater detail.

### Generation length and constancy

Constancy, typically assessed through the coefficient of variation (Sheeba *et al*. 1998; Prasad *et al*. 2003), or more recently through the fluctuation index (Dey & Joshi 2006; Dey *et al*. 2008, 2012; Pandey & Joshi 2026a,b), is the inverse of the extent of fluctuations in population size over generations. Earlier work had shown that FEJs had evolved greater constancy than the JBs (Prasad *et al*. 2003; Dey *et al*. 2008). This increased constancy was attributed (Prasad *et al*. 2003; Dey *et al*. 2008) primarily to evolved changes in FEJ life-history traits, such as lowered fecundity, body weight, and pre-adult survivorship, compared to the JBs (Prasad *et al*. 2000, 2001).

In our study, the 18 day cycle treatment in the LH food regime represents conditions closest to earlier studies (21-day generation length; LH food regime). Our results for this treatment combination align with earlier work, showing that FEJs exhibit significantly lower FI relative to JBs, which in turn implies higher constancy for the FEJs (Figure 1). However, this pattern did not generalize across other treatment combinations. When assayed on a 12 day cycle, FEJs did not differ in constancy stability from the JBs. This highlights the substantial influence of generation length on observed constancy.

This observed outcome is consistent with the known ecology of the FEJs. FEJs are routinely maintained in the laboratory on a 10-day discrete generation cycle. Forward selection (∼600 more generations of selection relative to the earlier studies examining FEJ dynamics) has driven a suite of correlated changes in life-history-related traits, such as lower fecundity, smaller body size, lower lipid content, and a shortened lifespan, relative to the JBs (Prasad *et al*. 2000, 2001; Prasad 2003; Modak 2009). When assayed at a generation length substantially longer than their native maintenance cycle, FEJ females would be much older than their optimal oviposition window and experience an age-related decline in fecundity. On average, a 5-day-old FEJ female lays ∼10 eggs, and this fecundity further decreases (often dropping to zero eggs) when measured on an 11-day-old female (see Figures 3 & S4 in Rao *et al*. 2025a).

Additionally, FEJs tend to have characteristically higher mortality rates compared to JBs (Ghosh & Joshi 2012; Mital 2019; personal observations), reflecting the constraints imposed by the response to extremely stringent selection pressures. During the dynamics assay, vials occasionally experience crowding due to the presence of a high number of larvae, adults, or both. FEJs are known to experience high levels of adult mortality at low and high adult densities, ranging between ∼25% to ∼40% mortality (Rao *et al*. 2025a). The higher mortality of FEJs is attributed to their generally poor performance under stressful conditions. FEJs typically have low survivorship when subjected to various stressors such as larval crowding, increased waste concentration in their food medium and adult starvation (Prasad 2003).

The higher adult mortality characteristic of FEJs, coupled with their low female fecundity, would, in turn, affect their population dynamics – when few eggs are laid, only a few adults eclose and survive to reproduce each generation. The surviving females would also exhibit low fecundity, resulting in small population sizes with very low levels of variation around the mean. This was also observed in our study, where the average population size maintained by the FEJs is ∼20 adults in the 18 day cycle regime and ∼35 adults in the 12 day cycle regime (Figure 3b). These effects could together contribute to the increased constancy stability of the FEJs observed in the 18 day cycle length treatment, as consistent with their autocorrelation patterns (Figure S4).

The relaxed selection populations CRFs and FRFs generally showed decreased constancy stability, particularly in the LH regime. A significantly higher FI of FRFs relative to the JBs and FEJs was observed in the 12 day cycle under the LH regime (Figure 1, see sub-section on relaxed selected populations for more details).

### Generation length and persistence

Unlike constancy, persistence did not significantly differ between the selection regimes in most treatment combinations, aligning with the results obtained from previous studies, which suggest that persistence need not evolve in parallel with constancy (Dey *et al*. 2008). However, the FEJs experienced significantly higher extinctions than the JBs in the 18 day cycle treatment, particularly in the LL regime (Figures 2 & S2), implying lower persistence relative to the JBs. This result differs from that of Dey *et al*. (2008), in which the persistence of JBs and FEJs did not significantly differ statistically, although JBs did show a slightly higher absolute probability of independent extinctions. One speculation for this reversal of patterns can be linked to the marked decline in FEJ fecundity over a further ∼600 generations of forward selection. Dey *et al*. (2008) reported the fecundity of FEJs to be ∼ 40 eggs per female, measured at the end of their dynamics assay (after ∼160 generations of selection). This is much higher than the average fecundity observed in FEJs at the time of the present study (∼10 eggs per female on average: Rao *et al*. 2025a), and therefore, could have contributed to the relatively lower extinction rates observed in the earlier study.

For each generation, by the 18^th^ day in the assay, FEJ fecundity would have further declined due to the increased age of the females, and this, coupled with the lower lifespan of FEJs relative to JBs (Modak 2009), would lead to a greater risk of extinction. For any given cycle length, in the LH regime, flies are provided with a live yeast paste for two days before the next generation is initiated. Yeast is known to boost female fecundity in *Drosophila* (Chippindale *et al*. 1993) and, thus, could result in a lower extinction probability in this treatment combination relative to the LL regime (Figure 2).

Interestingly, in the 12 day cycle length treatment, JBs showed the highest extinction probability among the four selection regimes (Figures 2 & S2). Although the difference is not statistically significant, it may have some biological significance, as it shows that JBs experience greater demographic instability when assayed at a generation length shorter than their maintenance regime. When reared at moderate larval densities, JBs have the largest body size and the highest fecundity among the four selection regimes (Rao *et al*. 2025a).

Their high fecundity on day 12 (see Rao *et al*. 2025a) may contribute to the increased risk of extinctions observed at this cycle length. Extinctions would be primarily driven by populations crashing due to excessively high larval or adult densities achieved in the assay. Therefore, population stability may not depend solely on evolutionary history, but also on the alignment (or misalignment) between life-history traits and the cycle length under which it is assessed. Such genotype-by-environment interactions are well known (Falconer, 1981) and quite often also observed in laboratory evolution experiments (highlighted in Rose *et al*. 1996).

### Relaxed-selection populations show a general convergence towards their pre-forward selection ancestral state, exceeding it for some attributes

The relaxed-selection populations (CRFs and FRFs) provide some insight into the role of the rapid development and early reproduction selection pressures in shaping population dynamics. In our study, the estimates of stability or demographic attributes measured for the FRFs and CRFs were, for the most part, closer to the JBs than the FEJs, indicating a clear impact of the relaxation of rapid development selection pressure (Figures 1 and 2). Interestingly, the relaxed populations maintained higher intrinsic growth rates (*r*) and equilibrium population sizes (*K*) than the JBs (Figures 3 & 4). The higher growth rates (Figure 4), coupled with their higher fecundity relative to FEJs (see Rao *et al*. 2025a), might also be expected to make these populations more prone to extinction. We observed that the higher *r* did correlate with greater fluctuations in the relaxed populations, whose FI, particularly in the LH food regime, was higher than the JBs and FEJs (Figure 1). The FRFs and CRFs also exhibited a stronger two-point cycle relative to both FEJs and JBs, as observed in the autocorrelation analyses (Figure S4). One would expect that in such cases, these populations would be more prone to extinction. However, this did not translate into a greater number of extinctions, as the FRFs and CRFs had the lowest extinction probability across most conditions (Figures 2 and S2), perhaps a result of their higher equilibrium and average population size (Figure 3). It is known that more than population size fluctuations being large, it is actually how often a population hits very low sizes that has a greater effect in reducing persistence (Hildenbrandt *et al*. 2006; Grimm & Wissel 2004; Dey & Joshi 2013).

Our results once again indicate that constancy and persistence can evolve independently, indeed in opposing directions (Figures 1 and 2). While earlier work showed that these two components need not co-evolve (Dey *et al*. 2008), our findings show that enhanced persistence can co-evolve with reduced constancy. A similar ecological rather than evolutionary effect was also seen earlier in the context of a change in the magnitude of immigration in a *Drosophila* experiment, increasing persistence while reducing constancy (Dey & Joshi 2013). This underscores the importance of assessing and comparing the stability of populations across multiple axes. Along the axis of constancy, the relaxed selection populations appear relatively unstable amongst all selection regimes. However, on the axis of persistence stability, they consistently show the lowest extinction probabilities, particularly in the LL food regime. Thus, conclusions about population stability can qualitatively differ, depending on which axis of stability is emphasized.

The lower extinction probabilities can be, in part, explained by analyzing the competition coefficient (Figure S3), which shows that FRFs and CRFs tend to have less negative ⍺ values, suggesting reduced sensitivity to an increase in density. Another line of evidence is that FRFs and CRFs have evolved to have a smaller adult body size relative to JBs when cultured at moderate larval density (see Figure 1 in Rao *et al*. 2025a), consistent with their maintaining higher equilibrium sizes (Figure 3). Recent work on adult crowding in *Drosophila* has shown that smaller flies can endure adult crowding better (Rao *et al*. 2025a, b), with more flies surviving the crowding period, as well as the surviving females having higher fecundity than those subjected to low adult density. This aspect may also be at play in our assay, particularly when small flies form high-density groups of adults, as seen in the cases of FRFs and CRFs (personal observations). According to Rao *et al*. (2025a), such groups typically exhibit very low mortality rates (<10%). This would imply that several individuals survive the crowding and can contribute to the next generation in the assay. While fecundity may be negatively affected by the timing of the next generation’s initiation, FRFs and CRFs show significantly higher fecundity than FEJs, which would minimize the risk of extinction due to demographic stochasticity. However, they do have considerably lower fecundity than the JBs; consequently, the egg density/larval density achieved in these cultures may not be as high as in the case of JBs, thereby reducing the risk of extinction due to a population crash.

### An empirical perspective on population stability

Recent work examining *Drosophila* population dynamics (Pandey & Joshi 2026a,b), and also theoretical work (Tung *et al*. 2019), suggests that simple population growth models may not always capture all aspects of life-history evolution that can impact population dynamics and stability. In particular, Pandey & Joshi (2026b) point out that a set of populations evolving higher growth rates than ancestral controls across a range of densities spanning across *K*, without a concomitant increase in *r* as well, cannot be accommodated even within three-parameter models like the theta-logistic or theta-Ricker. In the present study, though various sets of populations have higher realized growth rates than others across various densities (Figures 5, 6), their realized growth rates all come close to 1 as they approach their respective *K* values. Moreover, the CRFs and FRFs do seem to sustain higher mean realized growth rates in the low density bins compared to the JBs (Figures 5,6), even though the differences between them in *r* estimates are not significant in most treatment combinations (Figure 4). Thus, unlike the MCU populations, adapted to larval crowding at low food levels, used by Pandey & Joshi (2026a), our populations in this study will likely be amenable to their dynamics being captured by simple population growth models.

Pandey & Joshi (2026a,b) further suggested that it would be useful to consider an empirical approach to the issue of the evolution of stability attributes by focusing on how various life-history-related traits, and their sensitivities to density, evolve under different selection regimes and then linking those trait-specific changes to alterations in the return maps which ultimately determine dynamics and stability characteristics. This is conceptually similar to the agent-based modeling approach of Tung *et al*. (2019), which links dynamics and stability with the sensitivity of life-history-related traits to density, and provides far better fits to *Drosophila* population size time series under a variety of conditions than the use of simple population growth models. Our findings are consistent with this suggestion, as they indicate that when populations are subject to selection pressures not involving crowding *per se*, the evolved dynamics might remain explainable by simple population growth models, whereas the dynamics of populations adapted to crowding may not necessarily be captured by such simple models. Like Pandey & Joshi (2026a,b) and Tung *et al*. (2019), we believe that such an empirical approach to understanding the evolution of dynamics and stability attributes deserves further study, both theoretically and through experimental evolution.

### Conclusions

1. We show that the relaxed-selection populations (CRFs, FRFs) evolved to have enhanced demographic attributes relative to their ancestral controls (relatively higher *r*, higher *K,* and lower ⍺ than the JBs), indicating the potential evolution of greater larval competitive ability in these populations. Future experiments could aim to test this prediction directly.
2. Different measures of stability, such as constancy and persistence, can evolve to change in opposite directions over the course of selection, mirroring their inverse changes under different immigration treatments (Dey & Joshi 2013). This highlights the importance of assessing population stability across multiple axes.
3. Unlike some populations adapted to larval crowding (Pandey & Joshi 2026a,b), the dynamics of the FEJs, JBs, FRFs, and CRFs do seem to be amenable to being captured reasonably well by simple population growth models like the Ricker (1954) model. This lends strength to the suggestion that an empirical approach linking dynamics and the sensitivity of life-history-related traits to density via return xmaps may be helpful in understanding the evolution of population stability under varying selection regimes.
4. Finally, our results highlight the significance of generation length and its influence on population dynamics. We find that a relatively small change in generation length (amounting to a 6-day difference in our experimental design) in a single species can qualitatively alter several aspects of population stability. Harmonizing generation length in a population dynamics experiment with the usual generation length experienced by the different populations should be an essential aspect of experimental design.

