## Supplementary Figures and Tables for "Revisiting the evolution of population stability due to selection for rapid development and early reproduction in *Drosophila*: the role of generation length"

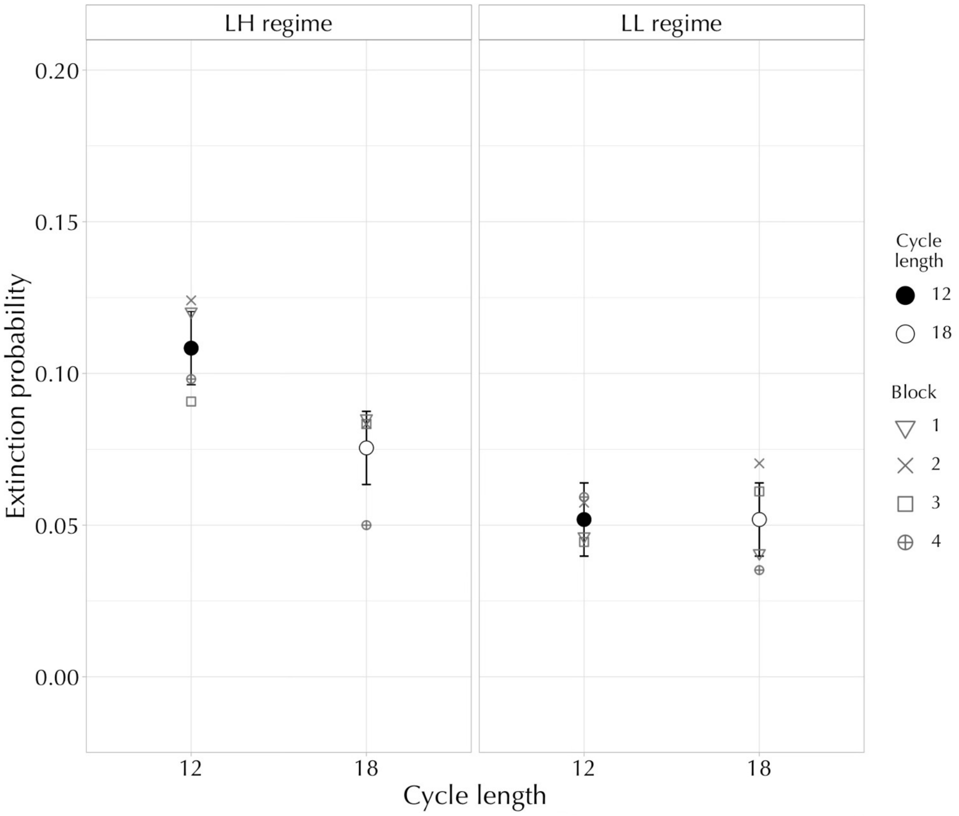


**Figure S1.** Mean extinction probability across combinations of two generation lengths and food regimes, averaged across other factors. Persistence is inversely related to extinction probability. The error bars represent 95% confidence intervals around the mean and can be used for visual hypothesis testing. The lack of an overlap between the error bars of any two groups signifies a statistically significant difference between their respective means.

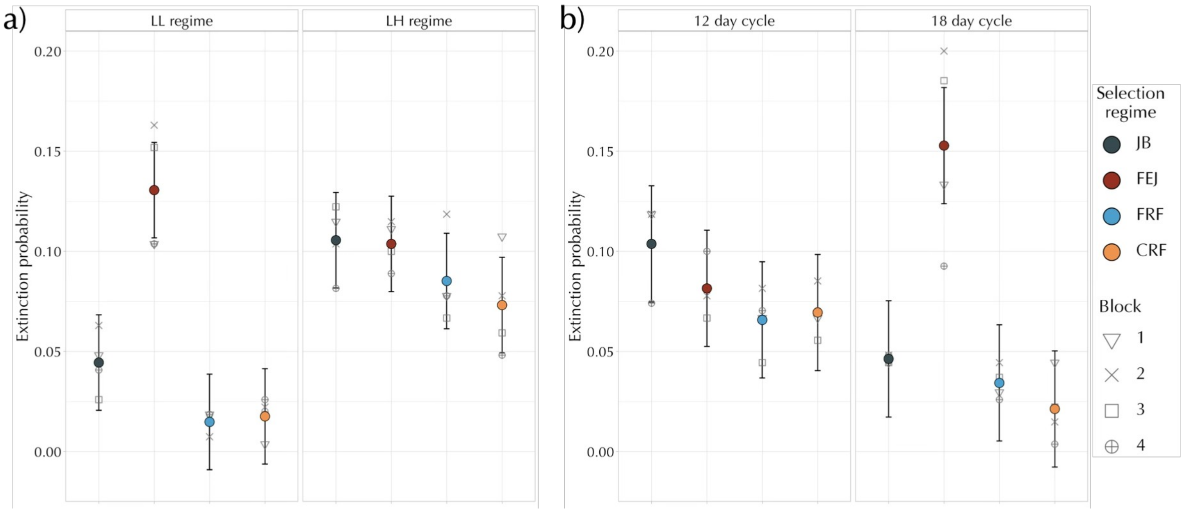


**Figure S2.** Mean extinction probability (per replicate population per generation) of the selection regimes across **a)** food regimes and **b)** generation length treatments, when averaged across other factors. Persistence is inversely related to extinction probability. The error bars represent 95% confidence intervals and can be used for visual hypothesis testing. The lack of an overlap between the error bars of any two groups signifies a statistically significant difference between their respective means.

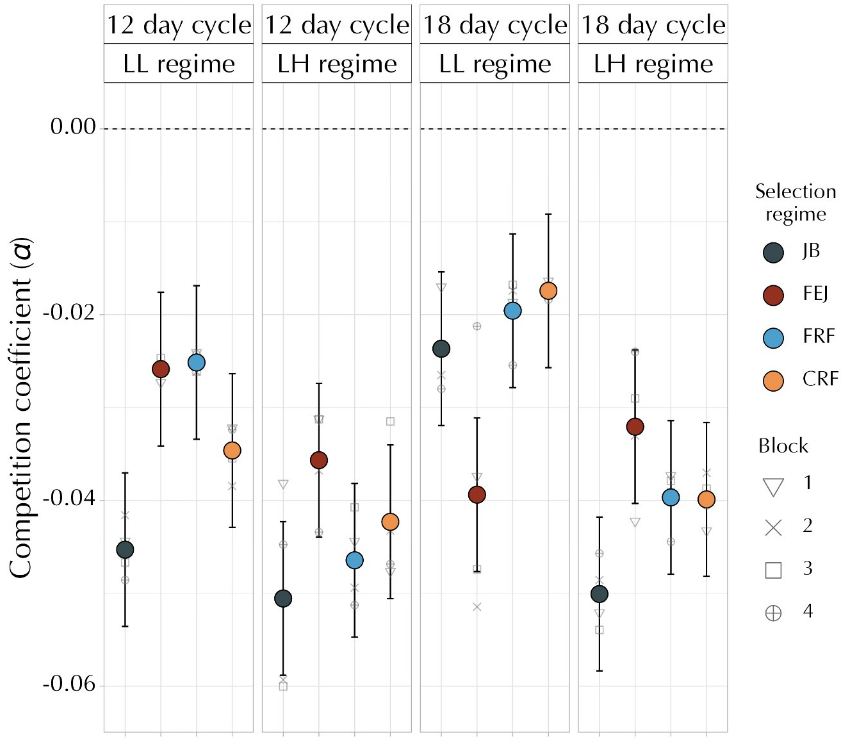


**Figure S3.** Estimates of the competition coefficient ($\alpha$) in the different selection regimes across all treatment combinations over the course of the population dynamics assay. The error bars represent 95% confidence intervals and can be used for visual hypothesis testing. The lack of an overlap between the error bars of any two groups signifies a statistically significant difference between their respective means.

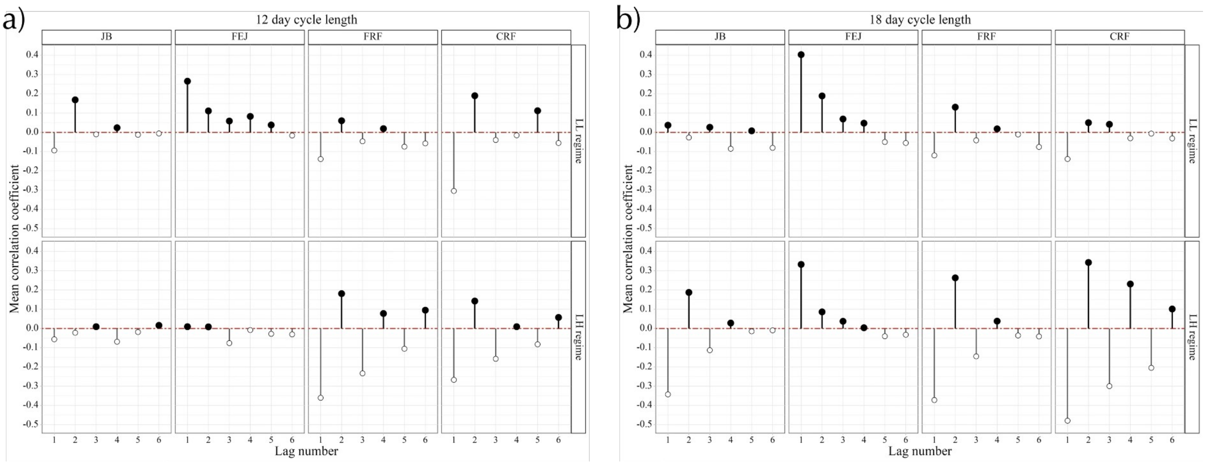


**Figure S4.** Mean autocorrelation coefficients of the selection regimes across both generation lengths and food regimes, averaged across replicate populations. The alternation of the positive and negative correlation coefficient signifies the presence of a two-point cycle. A stronger cycling between the two signs indicates a stronger two-point cycle in the population.

### Supplementary Tables

**Table S1.** ANOVA results for Fluctuation Index (FI). The effects of block cannot be estimated from our experimental design and are, therefore, omitted from the table.

| **Factor** | **df** | **MS** | ***F*** | ***P*** |
|---|---|---|---|---|
| Selection regime | 3 | 0.212 | 36.511 | < 0.001 |
| Day | 1 | 0.051 | 7.533 | 0.071 |
| Food regime | 1 | 2.476 | 603.702 | < 0.001 |
| Selection regime × Day | 3 | 0.046 | 3.657 | 0.057 |
| Selection regime × Food regime | 3 | 0.176 | 22.146 | < 0.001 |
| Day × Food regime | 1 | 0.096 | 19.419 | 0.022 |
| Selection regime × Day × Food regime | 3 | 0.101 | 18.102 | < 0.001 |

**Table S2.** ANOVA results for total extinction probability. In our design, the effects of block cannot be estimated and are therefore omitted from the table.

| **Factor** | **df** | **MS** | ***F*** | ***P*** |
|---|---|---|---|---|
| Selection regime | 3 | 0.017 | 49.368 | < 0.001 |
| Day | 1 | 0.004 | 3.800 | 0.146 |
| Food regime | 1 | 0.025 | 33.365 | 0.010 |
| Selection regime × Day | 3 | 0.014 | 10.640 | < 0.01 |
| Selection regime × Food regime | 3 | 0.008 | 9.020 | < 0.01 |
| Day × Food regime | 1 | 0.004 | 21.635 | 0.018 |
| Selection regime × Day × Food regime | 3 | 0.0003 | 1.710 | 0.233 |

**Table S3.** ANOVA results for estimates of the equilibrium population size (*K*). In our design, the effects of block cannot be estimated and are therefore omitted from the table.

| **Factor** | **df** | **MS** | ***F*** | ***P*** |
|---|---|---|---|---|
| Selection regime | 3 | 1925.6 | 59.40 | < 0.001 |
| Day | 1 | 630.509 | 34.22 | < 0.01 |
| Food regime | 1 | 3303.79 | 326.32 | < 0.001 |
| Selection regime × Day | 3 | 566.421 | 12.672 | < 0.01 |
| Selection regime × Food regime | 3 | 591.161 | 21.590 | < 0.001 |
| Day × Food regime | 1 | 465.659 | 22.752 | 0.017 |
| Selection regime × Day × Food regime | 3 | 165.251 | 11.969 | < 0.01 |

**Table S4.** ANOVA results for estimates of the intrinsic growth rate (*r*). In our design, the effects of block cannot be estimated and are therefore omitted from the table.

| **Factor** | **df** | **MS** | ***F*** | ***P*** |
|---|---|---|---|---|
| Selection regime | 3 | 1.674 | 63.261 | < 0.001 |
| Day | 1 | 0.336 | 24.726 | 0.015 |
| Food regime | 1 | 0.692 | 30.806 | 0.011 |
| Selection regime × Day | 3 | 0.108 | 4.064 | 0.044 |
| Selection regime × Food regime | 3 | 0.001 | 0.092 | 0.962 |
| Day × Food regime | 1 | 0.089 | 9.446 | 0.054 |
| Selection regime × Day × Food regime | 3 | 0.092 | 5.809 | 0.017 |

**Table S5.** ANOVA results for estimates of competition coefficient ($\alpha$). In our design, the effects of block cannot be estimated and are therefore omitted from the table.

| **Factor** | **df** | **MS** | ***F*** | ***P*** |
|---|---|---|---|---|
| Selection regime | 3 | 0.0003 | 8.602 | < 0.01 |
| Day | 1 | 0.0004 | 19.477 | 0.021 |
| Food regime | 1 | 0.0027 | 188.158 | < 0.001 |
| Selection regime × Day | 3 | 0.0002 | 4.672 | 0.031 |
| Selection regime × Food regime | 3 | 0.0002 | 6.865 | 0.010 |
| Day × Food regime | 1 | 7.75E-05 | 1.645 | 0.28 |
| Selection regime × Day × Food regime | 3 | 0.0002 | 10.869 | 0.002 |

**Table S6.** ANOVA results for the realized growth rate. In our design, the effects of block cannot be estimated and are therefore omitted from the table.

| **Effect** | **df** | **MS** | ***F*** | ***P*** |
|---|---|---|---|---|
| Selection regime | 3 | 46.989 | 26.96 | < 0.001 |
| Day | 1 | 0.081 | 0.091 | 0.78 |
| Food regime | 1 | 1.473 | 0.997 | 0.39 |
| Bin-size | 3 | 451.062 | 210.08 | < 0.001 |
| Selection regime × Day | 3 | 4.487 | 1.600 | 0.256 |
| Selection regime × Food regime | 3 | 1.948 | 1.176 | 0.37 |
| Day × Food regime | 1 | 0.473 | 0.326 | 0.60 |
| Selection regime × Bin-size | 9 | 16.011 | 10.01 | < 0.001 |
| Day × Bin-size | 3 | 2.970 | 1.945 | 0.19 |
| Food regime × Bin-size | 3 | 4.796 | 2.018 | 0.18 |
| Selection regime × Day × Food regime | 3 | 0.706 | 0.542 | 0.66 |
| Selection regime × Day × Bin-size | 9 | 0.895 | 0.339 | 0.95 |
| Selection regime × Food regime × Bin-size | 9 | 1.820 | 0.941 | 0.50 |
| Day × Food regime × Bin-size | 3 | 0.537 | 0.751 | 0.54 |
| Selection regime × Day × Food regime × Bin-size | 9 | 0.233 | 0.126 | 0.99 |
